# Bio-inspired chromatophore network model generates dynamic skin patterns akin to those observed in cephalopods

**DOI:** 10.64898/2026.09.30.755305

**Authors:** Yasemin Ersoy, Robin Peter, Gabriel Barello, Emily E. Meyer, Emily L. Mackevicius, Stephen L. Senft, Roger T. Hanlon, G. Bard Ermentrout, Stephanie E. Palmer

**Author notes:** Department of Radiation Oncology, University of California San Francisco, San Francisco, California, USA. Department of Biomedical Informatics, University of Colorado Anschutz, Aurora, Colorado, USA.

## Abstract

Cephalopod skin exhibits a wide range of dynamic visual displays, including traveling waves, light/dark flashes, and flickering, all generated by muscle-regulated expansion and retraction of pigment-filled elastic sacs known as chromatophores. While central neural control is essential for high-level body-wide pattern coordination, experimental evidence from excised and denervated skin reveals that intricate spatiotemporal patterns can also emerge spontaneously in the skin. This raises questions about the respective roles of central versus peripheral control. To determine the simplest control necessary to generate such patterns, we developed ChromatoNet, a biophysical model of chromatophore arrays that includes mechanical interactions on 1- and 2-dimensional lattices with the excitability properties of radial muscles. With minimal external input, our modified Morris–Lecar model reproduces a rich repertoire of skin dynamics, including spontaneous flickering as well as traveling and spiral waves. Both spontaneous and stimulus-evoked waves emerge at an intrinsic resonant frequency, determined by model parameters. Propagation speeds remain stable across conditions and are consistent with the range observed in denervated skin preparations. The model generates diverse output states in response to local couplings and the time-dependence and shape of the input stimulation, with little top-down direction. The results provide a proof-of-principle that chromatophore/muscle dynamics with minimal central neural driving can produce rich dynamic patterns.

**Author summary:** Cephalopods are capable of a wide variety of dynamic patterns for camouflage and communication, although much about how these effects are physically generated in the skin remains unknown. Cuttlefish and other cephalopods can alter their appearance with remarkable speed, producing body-wide changes in less than one second [12]. At a granular level this relies on thousands of pigment-filled saccules, “chromatophores” that expand and retract to generate shifting coordinated patterns that provide the optical illusion of movement across the skin. Each sacculus is actuated by 18-30 radial muscles whose orchestrated contraction pulls it out horizontally into a flat disc of visible pigment. While a rich variety of such coordinated changes in chromatophore actuation can be generated by electrical stimulation of the brain, and by active behavior, similar patterns also occur in excised and denervated skin, pointing to at least some role for purely local control. To investigate whether that role might be substantial, we built a simple model that includes core features of the known electrical activity of the muscles as well as experimentally motivated underlying geometrical connections that capture the way neighboring chromatophores may directly influence one another. Our system produces traveling waves, flickering, and other dynamic motifs observed in cephalopods in the wild, without requiring detailed instructions from a “brain.” Small changes to how the system is wired or perturbed lead to large shifts in modeled skin patterning and dynamics, suggesting that in nature, across species and even within an animal, the skin itself may be tuned to display a wide space of possible patterns.

## Introduction

Cephalopod skin is one of the most dynamic displays in the animal kingdom [13]. Squid, cuttlefish, and octopus coordinate thousands of distensible pigmented chromatophore organs into traveling waves, spirals, and body-wide flashes within fractions of a second, generating patterns used for camouflage and communication [12, 19, 22, 39]. Each chromatophore organ has a pigment sac surrounded by radial muscles; when the muscles contract, the sac is pulled into a flat disc that exposes thousands of small pigment granules in the animal’s skin (Fig 1A) [10]. Because of the complexity of these patterns coupled with the size constraints of the animal’s brain, understanding how control of these patterns is distributed between the brain versus the skin remains a central challenge in the study of cephalopod neurophysiology and behavior.

**Fig 1.**
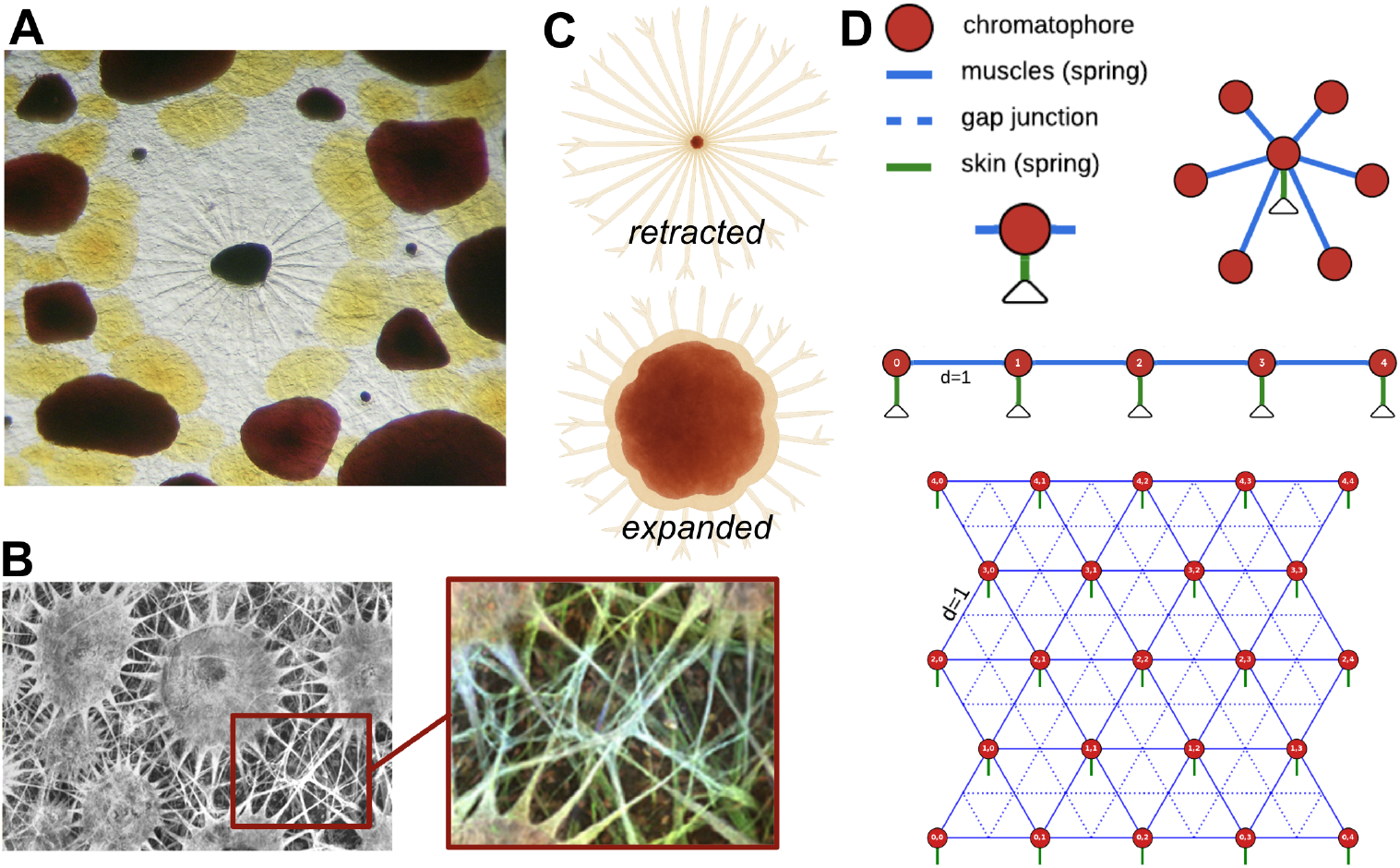
Chromatophore anatomy and model network architecture. (A) A close-up of the mantle skin of a squid, where the muscles of the semi-expanded brown chromatophore in the center are visible, surrounded by the lattice of expanded smaller yellow and red chromatophores. Three small punctate chromatophores are also evident. Image courtesy of Lydia M. Mäthger of the Hanlon Laboratory, used with permission. (B) Confocal image of squid chromatophores with a *β*-Actin antibody staining, illustrating the dense muscle network. Zoom in crop of the radial muscle fibers is pseudocolored by depth to make their three-dimensional layering clearer (adapted from Bell et al., 2013 [1]). (C) A cartoon of a single chromatophore unit in the retracted (top) and expanded (bottom) states, with radial muscles spanning the pigment sacculus. (D) Diagram of 1D and 2D hexagonal model networks. Red circles: chromatophores with variable size; blue lines: muscles (springs); blue dashed lines: gap junctions between muscles; green lines (attached to triangles): skin springs anchoring each chromatophore to its resting position. Chromatophore labels and indexing conventions are shown.

The historically prevailing framework attributes this coordination entirely to central neural control: circuits in the chromatophore lobes are known to monosynaptically drive each region of the skin, and lobe stimulation in higher motor centers of the brain (optic lobes, lateral basal lobes) evokes stereotyped displays [2, 6, 7, 12, 22, 44]. This architecture accounts for how the brain selects and initiates motifs, but each motorneuron controls only a local patch [7, 19, 22, 32]. It therefore remains unknown how wave-like patterns spread rapidly across the skin.

A growing body of observations points to an active role for the skin itself.

Anatomical work has revealed direct contacts between the radial muscles of neighboring chromatophores, providing a structural basis for local propagation [20, 31, 38] (Fig 1B). In excised and chronically denervated preparations, wave-like chromatophore activity, what Packard termed “wandering clouds,” emerges spontaneously without any descending neural input, and local electrical stimulation of excised skin reliably evokes both propagating and spiral waves [27, 28, 35]. The “passing clouds” of living cuttlefish, bands of expanded chromatophores that travel at a characteristic speed across an otherwise static background, illustrate this vividly in behavior [19, 28] (S1 Video). One parsimonious interpretation is that the brain modulates an active substrate rather than specifying wave dynamics element by element [24, 27].

We developed ChromatoNet to examine what spatiotemporal patterns a coupled, peripheral chromatophore network might produce on its own. We modeled each chromatophore as an excitable set of muscle elements governed by Morris–Lecar dynamics [23], linked to its neighbors through mechanical springs and gap junctions on a lattice, as shown in Fig 1. Central input can be applied locally and varied independently of the peripheral coupling. ChromatoNet reproduces spontaneous flickering, traveling waves, spirals, and pacemaker oscillations, with a small number of parameters controlling the transitions between behavioral regimes, consistent with the idea that the skin operates as an active dynamical system, with neuromodulatory input initiating displays rather than specifying each pattern *de novo*.

## Materials and methods

### Biological assumptions and model setup

The model operates on a single pigment layer of monochromatic chromatophores while multi-layer and multi-color extensions are considered in the Discussion. The model tests how much central control from the brain can be replaced with simple inputs and still produce complex patterns in the skin. The model rests on three assumptions, each drawing upon observations of chromatophore anatomy and physiology.

#### 1. Chromatophores expand and retract with muscle control

A single chromatophore organ consists of a pigment sacculus and its surrounding radial muscles, each anchored in the dermis(Fig 1C). Upon neural stimulation, muscle contraction pulls the punctate sacculus into a wide flat disc of pigmented color; muscle relaxation allows elastic recoil of the sacculus to close it [6, 10, 28]. We treat each chromatophore as a point whose effective size tracks the length of its attached muscles, and we model each muscle as an excitable active spring with a membrane voltage *V*_*µ*_ and potassium gating variable *W*_*µ*_ (Fig 1D). Each artificial chromatophore is also tethered to the surrounding skin by a passive restoring force, which we represent as a “skin spring”.

#### 2. Neighboring chromatophores are mechanically and electrically linked through their muscles

The radial muscles of nearby chromatophores can make direct contact with one another [20, 38], so a muscle contracting one chromatophore could stretch muscles of its neighbor (Fig 1B). Electrophysiological evidence suggests gap-junction coupling among muscle fibers acting on the same chromatophore [10, 20, 31]. In our model, muscles sharing a chromatophore attachment point interact both mechanically, through stretch-sensitive calcium channels, and electrically, through gap junctions (Fig 1D).

#### 3. Chromatophore activity can arise spontaneously or be triggered by electrical or mechanical perturbation

Wave-like chromatophore activity emerges spontaneously in excised and denervated live skin preparations [27, 28, 35], and chromatophore expansion can also be experimentally evoked by direct electrical stimulation of motor neurons or the surface of the skin, and by local mechanical vertical depression or lateral displacement of the skin [28, 35, 38]. The model accommodates all of these perturbation modes, although for simplicity most simulations in this paper use electrical input. Electrical input enters the voltage equation as a spatially and temporally specified applied current *I*_app,*µ*_; mechanical input is modeled as a direct positional displacement of one or more chromatophores from their rest positions.

### Network geometry

To explore how the simplest models on chains connect to fuller skin simulations, chromatophores are placed on a one-dimensional chain or on a two-dimensional hexagonal lattice. Their connections approximate the center-out arrangement of real chromatophores in the tightest packed nearest neighbor network, with *N* × *M* nodes (Fig 1D). The total chromatophore count for the 2D case is *M* ·*N* ™⌊*M/*2⌋ due to the offset-row geometry used to approximate the hexagonal configuration of chromatophores. Each chromatophore *i* has a fixed rest position **z**_*i*_ and a dynamic position **x**_*i*_(*t*). Nearest-neighbor chromatophores are spaced one unit apart at rest.

Radial muscles connect each chromatophore to its immediate neighbors: in the 1D case, each interior chromatophore has two muscle connections; in the 2D hexagonal case, up to six. Each muscle *µ* connects a pair of chromatophores (*α, β*)_*µ*_: the muscle’s endpoints **y**_*µ*,*α*_ and **y**_*µ*,*β*_ track its attachments as the network deforms.

Gap junctions are placed between centers of neighboring muscles; in the 2D hexagonal lattice these connections form bow-tie patterns (Fig 1D). Boundary chromatophores (the endpoints of the 1D chain and the perimeter of the 2D lattice) are held fixed at their rest positions throughout each simulation, representing anchoring of the chromatophore layer to the broader tissue.

#### Spatial noise

Frozen Gaussian noise can be added to the resting positions of the chromatophores to represent static spatial variation in the chromatophore placement:

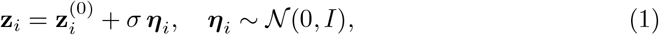

where 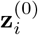 is the unperturbed lattice position and ***η***_*i*_ is sampled once at simulation initialization and held fixed thereafter.

#### Random connection removal

Real cephalopod skin is not a fully connected network of chromatophores and muscles, as biological variation creates variability and defects in the network. To explore this in the model, a fraction *f* of the network’s muscle connections is selected uniformly at random without replacement and removed from the network, along with the gap junctions associated with those muscles. Each removal fraction [0%, 10%, 20%, 30%, 40%, 50%, 70%, 100%] was tested across 100 independent random seeds.

### Model Dynamics

#### Chromatophore positions and muscle endpoints

Each chromatophore position **x**_*i*_ is governed by the skin spring force tethering it to its rest position **z**_*i*_ through *k*_*s*_, and the total pull from its attached muscle neighbors at locations **y**_*µ*,*i*_ via *k*_*c*_:

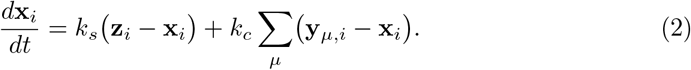

Each muscle attachment point pulls its chromatophore along the muscle axis by combined passive and active spring forces

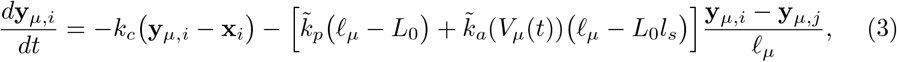

where *l*_*µ*_ = ∥**y**_*µ*,*i*_ − **y**_*µ*,*j*_∥ is the current muscle length, *L*_0_ is its resting length, and *l*_*s*_ *<* 1 is the fraction of the contracted length that creates the resultant length, *l*_*s*_ · *L*_0_. The active spring constant is voltage-gated, *k*_*a*_(*V*_*µ*_) = *k*_max_ · *σ*(*V*_*µ*_; *V*_th_, *V*_shp_), where *σ*(*v*; *θ, s*) = 1*/*(1 + *e*^*™*(*v™θ*)*/s*^) is a sigmoid function. To capture the experimentally observed asymmetry between rapid expansion and slower retraction of the chromatophores caused by their surrounding muscle activity [1], passive and active stiffnesses 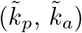 are reduced when a muscle is lengthening past its rest length, 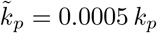 and 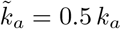 *k*_*a*_ when *l*_*µ*_ ≥ *L*_0_.

#### Membrane voltage and recovery

Each muscle’s membrane voltage *V*_*µ*_(*t*) follows Morris–Lecar dynamics [23] with dimensionless parameters and variables [33]:

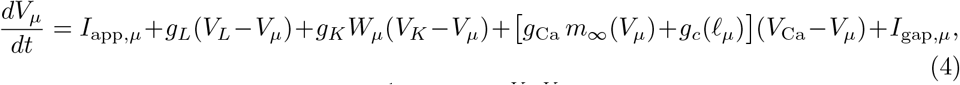

with calcium activation 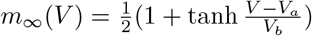. The stretch-sensitive calcium conductance channel opens when a muscle is elongated beyond threshold, providing the mechanochemical coupling through which stretch on one muscle can trigger voltage changes on its neighbors 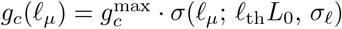. The recovery variable *W*_*µ*_ evolves as

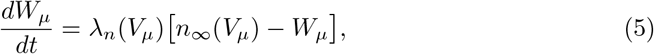

with 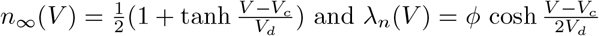.

#### Gap junction coupling

Neighboring muscles are electrically coupled through gap junctions (Fig 1D)

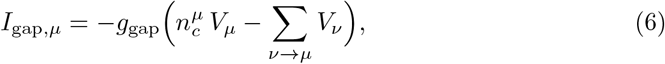

where the sum is over muscles *v* attached to the same shared chromatophore, *µ*, and *n*^*µ*^ counts these neighbors. Setting *g*_gap_ = 0 isolates purely mechanochemical propagation through *g*_*c*_.

### Numerical integration

Equations (2)–(5) are integrated using fourth-order Runge–Kutta. The simulation is implemented in PyTorch and runs on standard CPU or Apple Silicon hardware; a PyTensor implementation is available upon request. At high *k*_*c*_ values (*k*_*c*_ ≳ 2.6), the chromatophore-muscle coupling produces steeper voltage gradients that require finer time steps for stability (S1 Appendix). The simulation code and analysis scripts are available on GitHub at https://github.com/sepalmer/ChromatoNet.

### Parameters

Default parameter values are listed in Table 1. Voltage parameters follow standard Morris–Lecar values [23]; mechanical parameters were chosen to produce spatiotemporal patterns similar to those observed in cephalopod skin.

**Table 1.** Default model parameters. Voltage quantities are dimensionless (normalized to *V*_Ca_). Time is in seconds.

| Parameter | Symbol | Default | Description |
| --- | --- | --- | --- |
| Applied current | $I_{app}$ | 0.15 | Baseline stimulus current |
| Leak conductance | $g_L$ | 0.42 | Sets resting excitability |
| Leak reversal | $V_L$ | -0.5 | Leak reversal potential |
| K conductance | $g_K$ | 2.0 | Potassium channel conductance |
| K reversal | $V_K$ | -0.7 | Potassium reversal potential |
| Ca conductance | $g_{Ca}$ | 1.1 | Calcium channel conductance |
| Ca reversal | $V_{Ca}$ | 1.0 | Calcium reversal potential |
| Max stretch conductance | $g_c^{max}$ | 0.1 | Mechanochemical coupling strength |
| Stretch threshold | $\ell_{th}$ | 1.1 | Stretch activation threshold ($L_0$ units) |
| Stretch slope | $\sigma_\ell$ | 0.1 | Stretch activation sharpness |
| Gap junction conductance | $g_{gap}$ | 0.005 | Electrical coupling strength |
| Passive spring | $k_p$ | 0.1 | Muscle passive stiffness |
| Max active spring | $k_{max}$ | 2.0 | Voltage-activated muscle stiffness |
| Contracted length | $l_s$ | 0.33 | Contracted length as fraction of $L_0$ |
| Resting length | $L_0$ | 1.0 | Muscle resting length |
| Skin spring | $k_s$ | 0.5 | Tissue restoring force |
| Chromat.-muscle coupling | $k_c$ | 2.0 | Pigment sac-to-muscle attachment |
| Ca activation midpoint | $V_a$ | -0.01 | $m_\infty$ midpoint |
| Ca activation slope | $V_b$ | 0.15 | $m_\infty$ slope |
| K activation midpoint | $V_c$ | 0.0 | $n_\infty$ midpoint |
| K activation slope | $V_d$ | 0.3 | $n_\infty$ slope |
| Rate scaling | $\phi$ | 0.05 | Gating time constant scaling |
| Active threshold | $V_{th}$ | 0.0 | Active spring activation midpoint |
| Active slope | $V_{shp}$ | 0.005 | Active spring activation sharpness |
| Integration timestep | $\Delta t$ | 0.1 s | RK4 step (tunable per experiment) |
| Initial voltage | $V_{init}$ | -0.266 | Initial membrane voltage |
| Initial gating | $W_{init}$ | 0.144 | Initial gating variable |

## Results

### Generating waves

As a first test of ChromatoNet, we asked whether the network can produce propagating waves from a single, localized perturbation, as observed experimentally in denervated and excised cephalopod skin [26, 35, 38] (Fig 2B; S2 Video). We started with the 1D chain and then extended to the 2D hexagonal lattice.

**Fig 2.**
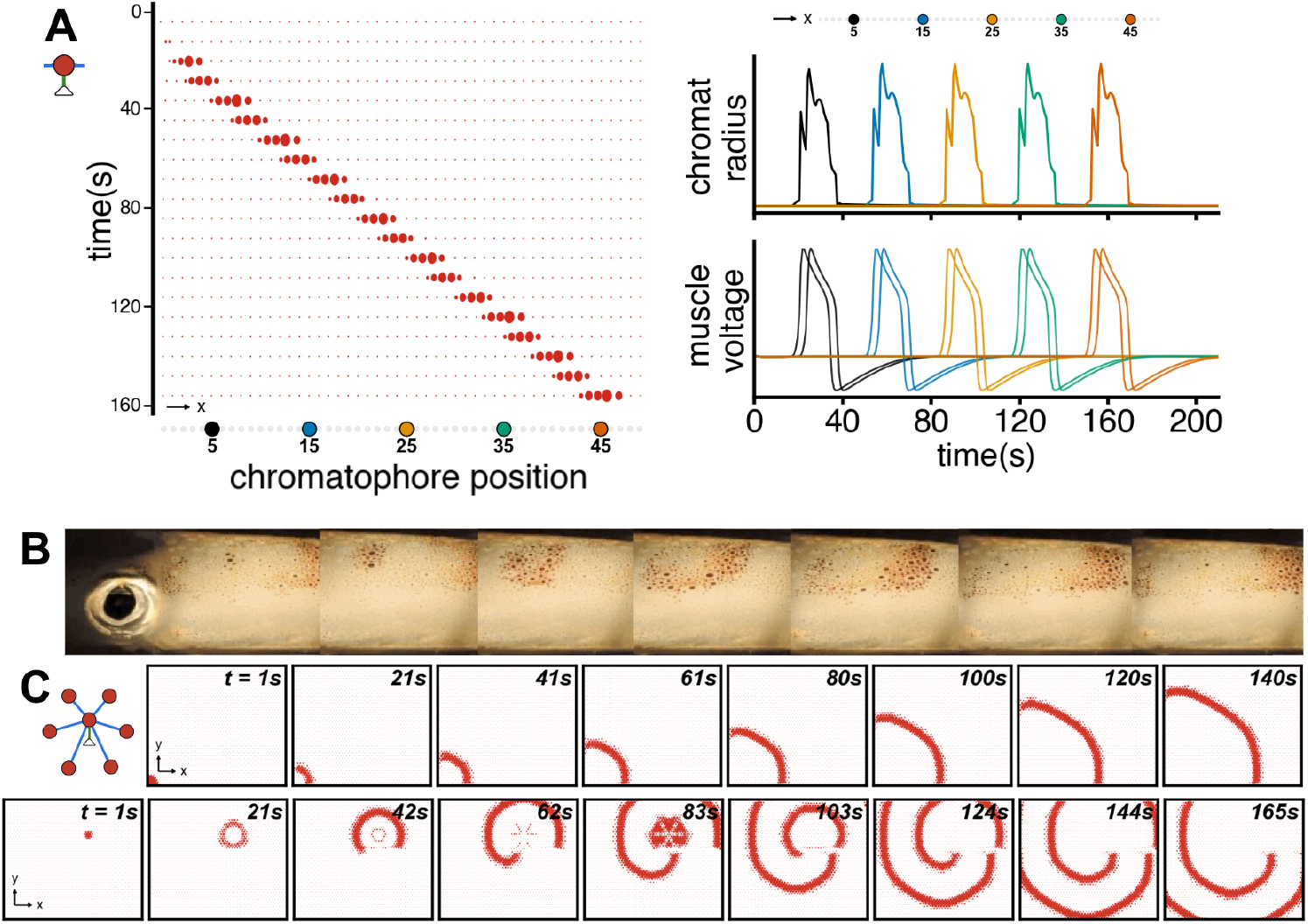
Peripheral dynamics generate traveling and spiral waves that resemble spontaneous cephalopod skin activity. (A) Traveling wave in the 1D model, initiated by a brief positional displacement of the leftmost chromatophore. Left: space–time snapshots of chromatophore positions along the chain. Right: colors 7/25 correspond to positions along the chain. Top: radius of five labeled chromatophores overtime; the sequential peaks confirm directed wave propagation. Bottom: membrane

In 1D, both positional and electrical stimulation can evoke a traveling wave. A positional displacement applied to the leftmost element of the chain produce a directed traveling wave: each chromatophore opens in sequence, with evenly spaced peaks in the radius traces show in (Fig 2A; S3 Video). Membrane voltage traces confirm that mechanical stretch initiates depolarization in each successive element with a characteristic refractory period, and an electrical stimulus to the same end of the chain produces an equivalent propagating wave through direct depolarization.

In the 2D hexagonal network, an electrical stimulus applied at a single corner produces an expanding wavefront that propagates coherently across the lattice (Fig 2C, top; S4 Video shows a comparable wave on a larger 50×50 lattice), recapitulating the body-wide passing-wave activity observed spontaneously or under direct electrical stimulation in denervated cephalopod skin [35, 38]. Placing the stimulus off-center near a rigid feature breaks the wavefront symmetry and seeds a persistent rotating spiral (Fig 2C, bottom; S5 Video). Spirals are a canonical regime of excitable media [41], and rotating spiral waves have been observed in denervated skin [26, 28], where a spiral emerges due solely to peripheral dynamics.

Once initiated, both the traveling wave and the spiral persist without continuous central input. The remainder of the Results characterizes how stimulus and parameter choices control which patterns the network produces.

### Varying Inputs

#### Stimulus shape and timing

Denervated cephalopod skin, including that of the squid, produces traveling and reentrant waves in response to simple electrical or physical stimulation [9, 26, 35, 38]. Intact animals, generally excepting squids, incorporate periodic traveling patterns, like passing clouds, into camouflage and communication displays [12, 14, 15, 19]. In ChromatoNet, setting the initial voltage of a small region to a spatially Gaussian depolarized state is enough to drive a single wave across the lattice without distortion (Fig 3A, top). Changing the geometry of the input into an asymmetric narrow strip generates a wave, that re-enters from above and below, leading to a self-sustaining periodic traveling wave (Fig 3A, bottom; S6 Video).

**Fig 3.**
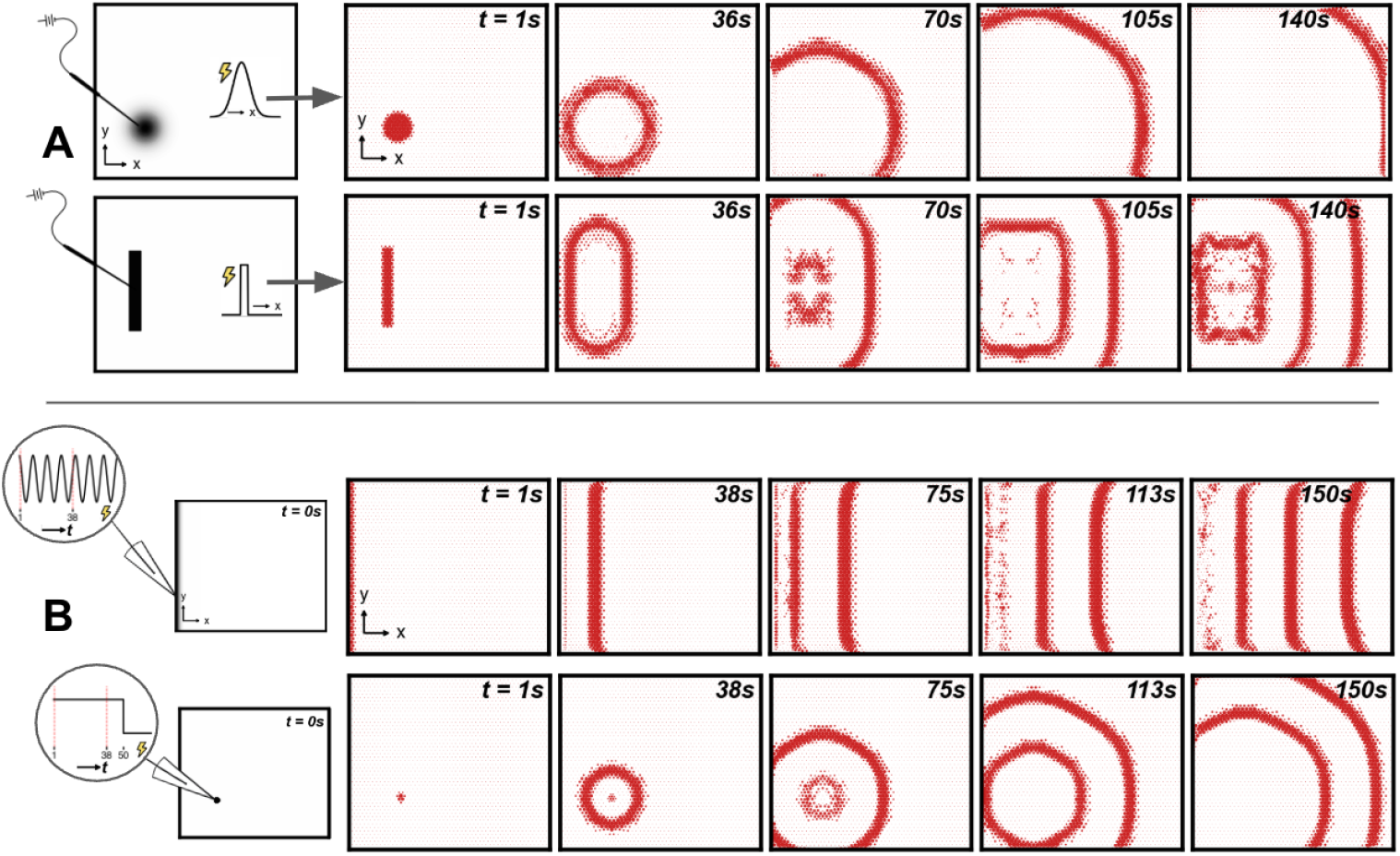
Stimulus shape and timing determine wave geometry. (A) Left: stimulus schematic showing a spatially broad Gaussian electrical perturbation applied near the lower-left of the chromatophore network (top) or asymmetric stimulation restricted to a narrow strip of the edge (bottom). Right: snapshots of the model chromatophores as waves propagate. A symmetric Gaussian stimulus (top) produces a single wavefront that propagates uniformly across the skin regardless of where it is applied. Breaking the spatial symmetry of the initial stimulation (bottom) causes the wave to reflect and reenter, generating a periodic traveling wave from a single, transient input. (B) Periodic waves with time dependent stimulation (left) with snapshots (right) of wave propagation. Top: periodic applied current along the left edge (sinusoidal temporal waveform shown). Bottom: sustained sub-threshold current applied at a small cluster of chromatophore muscles, showing periodic concentric rings emanating from the stimulation site, locked to the system’s intrinsic resonant frequency (∼20 mHz).

We explored two additional, time-varying, input types which produce periodic waves through different mechanisms. A continuous sinusoidal current generates a periodic train of waves (Fig 3B, top). A sustained sub-threshold current applied to a small interior cluster of chromatophore muscles produces repeated outward-spreading rings for as long as the current was held, with each new ring separated from the last relative to the network’s refractory period (Fig 3B, bottom). Constant input producing periodic output is consistent with focal expansion waves observed in electrically stimulated denervated squid skin [35]. The duration required to initiate a wave depends on system parameters. For certain settings, if the current is not held long enough, the wave cannot form; the low-pass filtering properties of the gap-junction-coupled muscle network can attenuate short inputs before they recruit neighboring chromatophores [20]. A localized command, even one that is constant in time, can act as a pacemaker for body-wide patterns.

#### An intrinsic frequency

Behavioral analyses of cuttlefish passing clouds have reported that waves maintain consistent spatial structure across a range of periods, suggesting a system-wide lock to a specific frequency [19]. The observation that constant input produced periodic output in ChromatoNet (Fig 3B, bottom) suggests the network has a characteristic frequency. We swept the periodic input frequency along the edge and measured the mean chromatophore opening frequency of the generated periodic plane waves: the output tracks the input only within a narrow band; outside that band, output settles near 20 mHz (Fig 4A). The network exhibits bandpass behavior, with a preferred operating range that pulls activity toward it.

**Fig 4.**
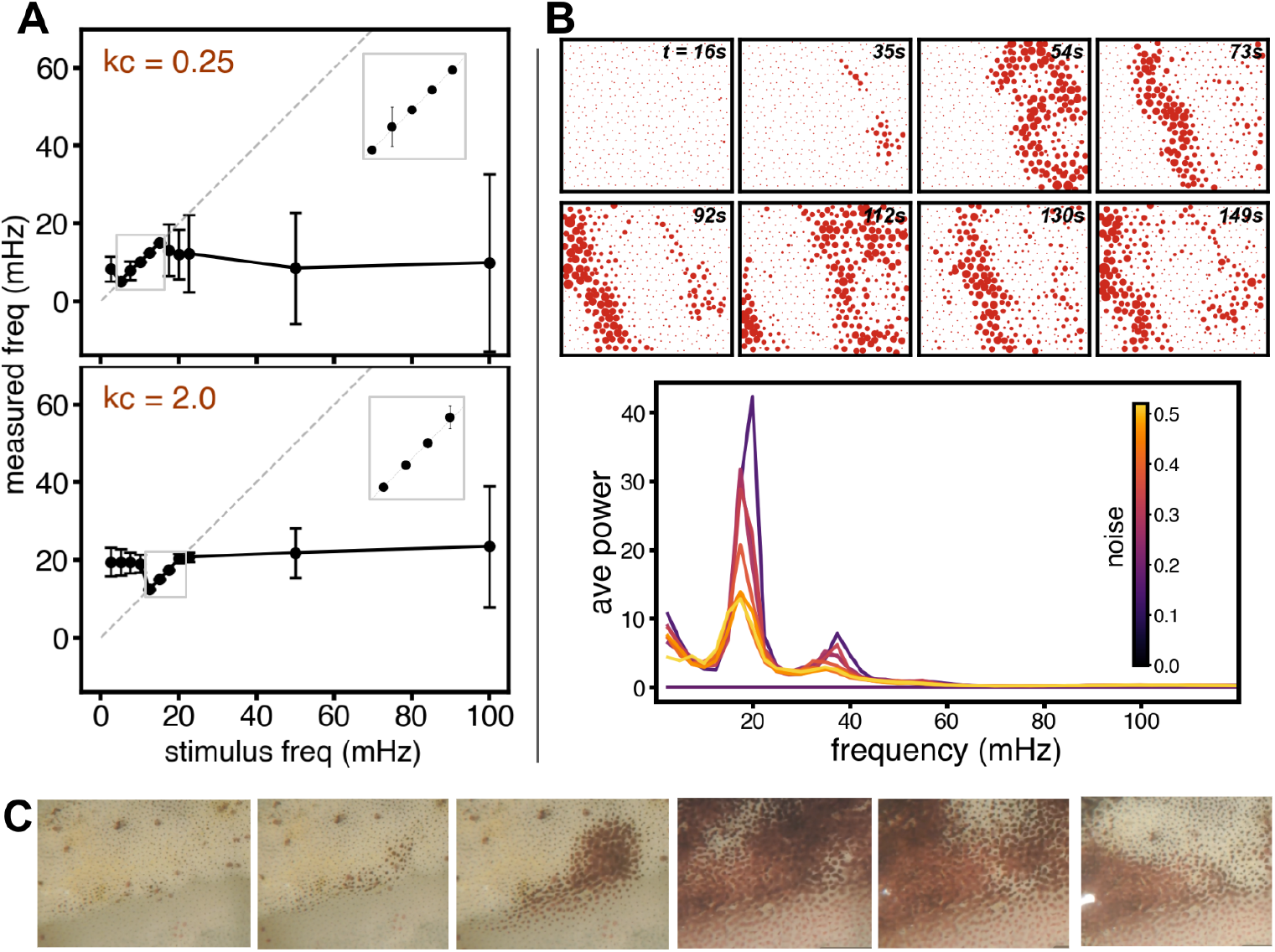
2D model input/output properties: intrinsic frequency even in spontaneous wave generation. (A) Measured traveling wave frequency as a function of the periodic input stimulus frequency, for two values of the chromatophore–muscle coupling constant *k*_*c*_ (*k*_*c*_ = 0.25, top; 2.0, bottom). Points show mean± 1 SD across interior chromatophores grouped by *x*-position. The dashed unity line indicates perfect input–output frequency matching. For both parameter regimes the output frequency tracks the input within a band near the system’s intrinsic frequency; outside this band, output locks to the intrinsic frequency, demonstrating bandpass filter behavior. Increasing *k*_*c*_ shifts the resonant frequency slightly upward. (B) Top: spontaneous activity driven by spatial Gaussian noise (*σ* = 0.2) added to the resting positions of the chromatophores. Snapshots of the 2D network at eight time points show stochastic wave fragments that nucleate, propagate, and dissolve without any external stimulus. Bottom: power spectral density of chromatophore sizes averaged across the network, for noise amplitudes ranging from *σ* = 0.00 to 0.52. A sharp peak near 20 mHz dominates at intermediate noise levels (0.20–0.32). At high noise amplitudes (*σ* ≳ 0.5) the spectral peak broadens as activity becomes increasingly incoherent. (C) Close-up snapshots over time of continuous spontaneous activity in denervated and pinned *Ascarosepion pfefferi* skin. Video courtesy of the Hanlon Laboratory.

This property is also reflected in wave speed: a wave’s spatial period and temporal frequency are tied through its velocity, so a system with a preferred frequency has a preferred wave speed. ChromatoNet waves travel at a consistent velocity for given mechanochemical parameters, and across denervated and excised cephalopod skin, propagation velocity is similarly stable across observations, falling near ∼ 1 cm/s [19, 26, 35, 38].

This intrinsic frequency is tunable. Increasing the chromatophore–muscle coupling constant *k*_*c*_ shifts the preferred frequency upward (Fig 4A, bottom), exposing a biophysical dial that tunes the skin’s operating point between regimes without altering the qualitative character of wave propagation.

#### Spontaneous waves and flickering

Excised and denervated cephalopod skin can produce wave activity without any stimulation [26, 35, 38](Fig 4C; S7 Video), and persistent flickering is a hallmark of the behavioral chromatophore display in some species, such as (*Dosidicus gigas* [34]. We add static Gaussian noise to the resting positions of the chromatophores in ChromatoNet, which could represent natural variation in chromatophore spacing across the skin, and are thereby able to produce spontaneous wave fragments that travel across the lattice (Fig 4B, top; S8 Video).

Generating spontaneous activity requires spatial noise above a threshold (*σ*≈0.2); below it the system is sub-excitable and fluctuations dissipated before recruiting neighbors. The threshold depends on the gap junction conductance (set to 0.005 in the figures).

Once noise exceeds the excitation threshold, spectral analysis reveals a sharp peak near 20 mHz at every noise amplitude (Fig 4B, bottom), the same frequency the system locks to under asymmetric initial voltage perturbations, driven periodic, and constant localized inputs. Spontaneous activity takes different forms depending on noise level: just above threshold, the power peak is tall and narrow and the network produces cleaner traveling waves; at higher noise, the peak broadens and activity resembles flickering, with shorter, less coherent fragments.

Both regimes remain active without central drive. Persistent flickering can therefore be sustained as a low-effort behavior of the peripheral network, under appropriate coupling conditions, without actively driving each “twinkle”.

### Varying parameters

#### Global parameters tune wave properties

Cephalopod skin tissue varies across species, body regions, and developmental stages. Muscle-to-muscle network density and chromatophore density both vary across the body, and behavioral repertoires range from static camouflage to high-frequency communication flashing [27, 28, 34, 35, 38]. We examine how varying biologically interpretable parameters of ChromatoNet shape wave dynamics, and we characterize the roles of five key parameters below. For each sweep, an electrical pulse is applied along the entire left edge of the network to create a wave propagating rightward.

The leak conductance, *g*_*L*_, sets baseline muscle excitability by determining how quickly voltage decays toward rest. This parameter governs the balance between traveling waves and full-skin flashes. At high *g*_*L*_ individual muscles require a stronger stimulus to excite; the wave slows and rounds at the edges. At low *g*_*L*_ the network becomes too excitable to generate a well-defined and spatially localized wave, and all chromatophores open nearly simultaneously in a full network flash (Fig 5A, top), like the body-wide synchronous flashing of *Dosidicus gigas* [34].

**Fig 5.**
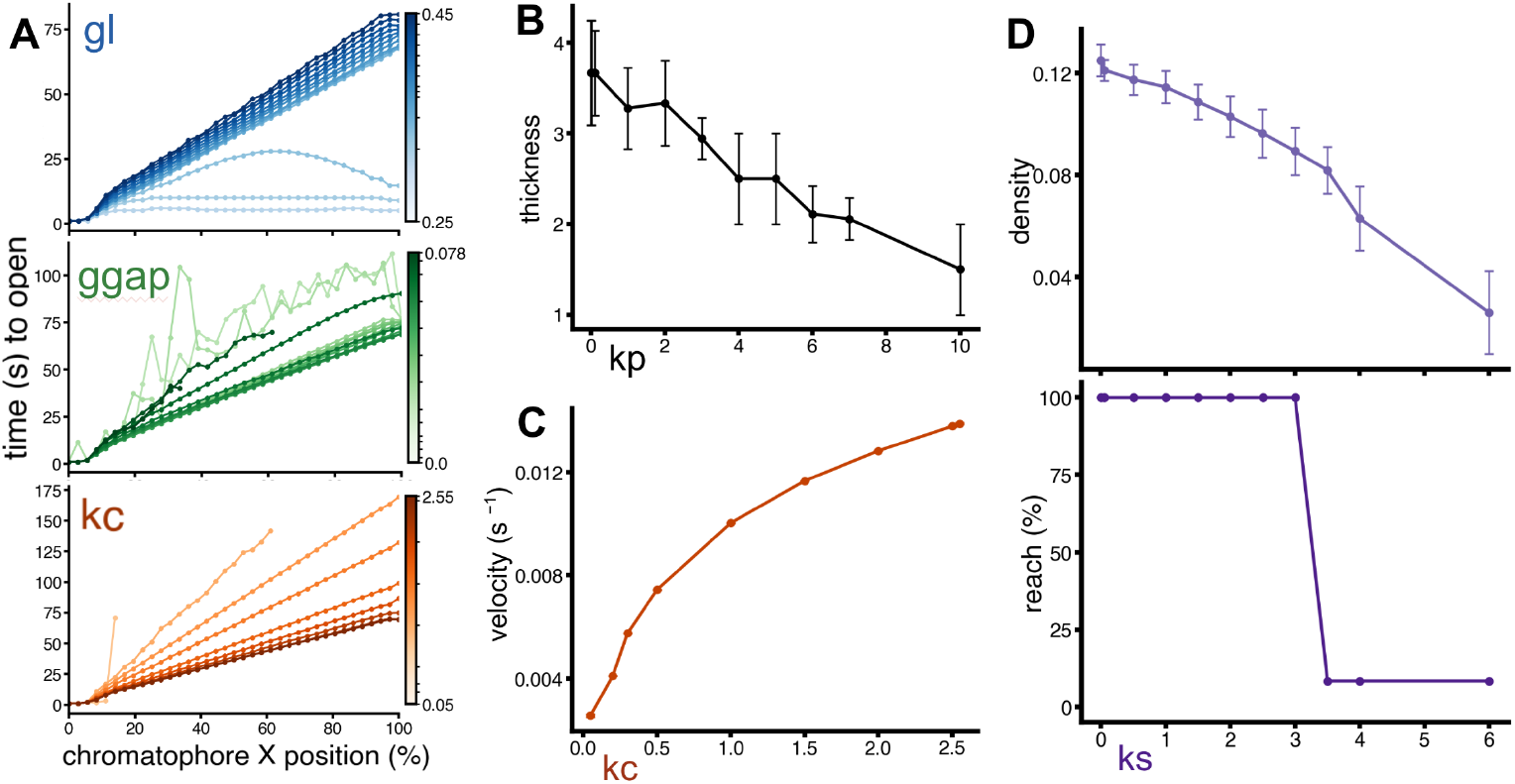
Global model parameters tune distinct properties of traveling waves. (A) Each panel shows wave propagation profiles (time for each chromatophore column to first reach 90% of peak radius, plotted against *x*-position) across a range of global parameter values. Top: leak conductance *g*_*L*_ (blue, *g*_*L*_ = 0.25–0.45). Higher *g*_*L*_ increases membrane shunting, slowing the wave and rounding its spatial profile. At low *g*_*L*_ the network approaches an all-or-none “flash” regime in which chromatophores across the entire skin expand nearly simultaneously. Middle: gap junction conductance *g*_gap_ (green, *g*_gap_ = 0.00–0.078). At low *g*_gap_, the wave is driven primarily by mechanical stretch and is sensitive to positional noise, causing dispersion. Moderate *g*_gap_ sharpens the wave front and increases propagation speed. Above a critical value (*g*_gap_ ≳ 0.08) gap junctions smooth voltage gradients too strongly and the wave fails to propagate. Bottom: chromatophore–muscle coupling *k*_*c*_ (orange, *k*_*c*_ = 0.05–2.55). Lower *k*_*c*_ causes the velocity to be so low that the wave cannot propagate, while increasing *k*_*c*_ monotonically increases wave velocity. (B) Wave thickness decreases as *k*_*p*_ increases (black). (C) Velocity increases as *k*_*c*_ increases (orange). (D) Open chromatophore density as a function of *k*_*s*_ (purple) and the fraction of the network reached by the wave (propagation reach) shows that increasing *k*_*s*_ progressively reduces both wave density and wave reach until propagation fails above a critical stiffness.

The gap junction conductance, *g*_gap_, represents the strength of electrical coupling between neighboring muscles. We find that *g*_gap_ acts non-monotonically on wave propagation. With *g*_gap_ at zero, propagation is carried entirely by mechanical stretch and the wave disperses in all directions rather than carrying a coherent wavefront. At moderate values, the gap-junctions couplings synchronize neighboring muscle voltages, sharpens the wavefront, and increases propagation speed. At sufficiently high values, the same coupling that supported propagation begins to oppose it: voltages equilibrate across the network faster than the mechanochemical signal can build a gradient, and the wave dies off (Fig 5A, middle). Electrical coupling conductance has an intermediate optimum: under-coupling distorts and disperses the wave, while over-coupling smooths the gradients that wave requires in order to propagate.

The chromatophore-muscle coupling, *k*_*c*_, sets how strongly muscle contraction stretches the chromatophore it pulls on. *k*_*c*_ heavily affects wave speed, and velocity is positively related to *k*_*c*_ (Fig 5A, bottom; Fig 5C). As *k*_*c*_ falls below a certain threshold, the wave moves so slowly that it can no longer excite its neighbors and fails to reach the end of the lattice, with slower waves traveling even less far.

Passive muscle stiffness, *k*_*p*_, determines how strongly each muscle resists stretch in its relaxed state. Increasing *k*_*p*_ produces thinner wavefronts with fewer simultaneously-open chromatophores (Fig 5B). The thinning occurs because the muscles relax back to their resting positions more aggressively as the restoring force is increased.

The skin spring constant, *k*_*s*_, relaxes each chromatophore back toward its rest position. Increasing *k*_*s*_ reduces both the distance a wave travels and the intensity with which the chromatophores opened along the way, appearing as a less densely colored pattern (Fig 5D, top; S1 Fig). Above *k*_*s*_ ≈ 3.5 the chromatophores barely open and the restoring force overwhelms the mechanical stretch signal, so waves fail to travel (Fig 5D, bottom).

These five parameters act on different pattern features and each has a plausible biological correlate, providing a biophysical framework where a small number of mechanisms might efficiently reshape wave dynamics across behavioral contexts; we develop these correspondences in the Discussion.

#### Rigid regions reroute waves

Real cephalopod skin is mechanically heterogeneous, with regions of higher stiffness and underlying musculature varying across the body surface [38]. Behavioral observations of passing clouds in cuttlefish show that traveling waves can navigate through and around regions of skin that simultaneously display static patterns, handling local mechanical heterogeneities without disruption [19, 27] (S1 Video). We created rigid regions in ChromatoNet by increasing the passive stiffness *k*_*p*_ and skin constant *k*_*s*_ within defined zones. Chromatophores in these stiff patches became too tightly bound to deform through neighboring activity. No new mechanism was needed; the same parameters that tune wave properties globally (Fig 5) can generate rigid heterogeneity when applied locally.

We first tested whether propagating waves can travel around rigid regions. A circular rigid region was placed in the upper right of the lattice with a small stimulus offset adjacent to it. A beautiful display of reentrant waves with a rigid region can be created by placing a circular rigid region in the upper right of the lattice, with a small symmetric stimulus offset, adjacent to the region. The wavefront is forced to split upon contacting the rigid region and the proximity causes asymmetry in the stimulus and creates a reentrant wave. The wave travels freely around the rigid region and a periodic pattern across the full lattice is established (Fig 6A; S9 Video). Stiff features in the network’s path can seed persistent spiral activity as well (Fig 2C, bottom), another example showing that the values of local parameters in the lattice contribute to pattern variations beyond simple obstruction.

**Fig 6.**
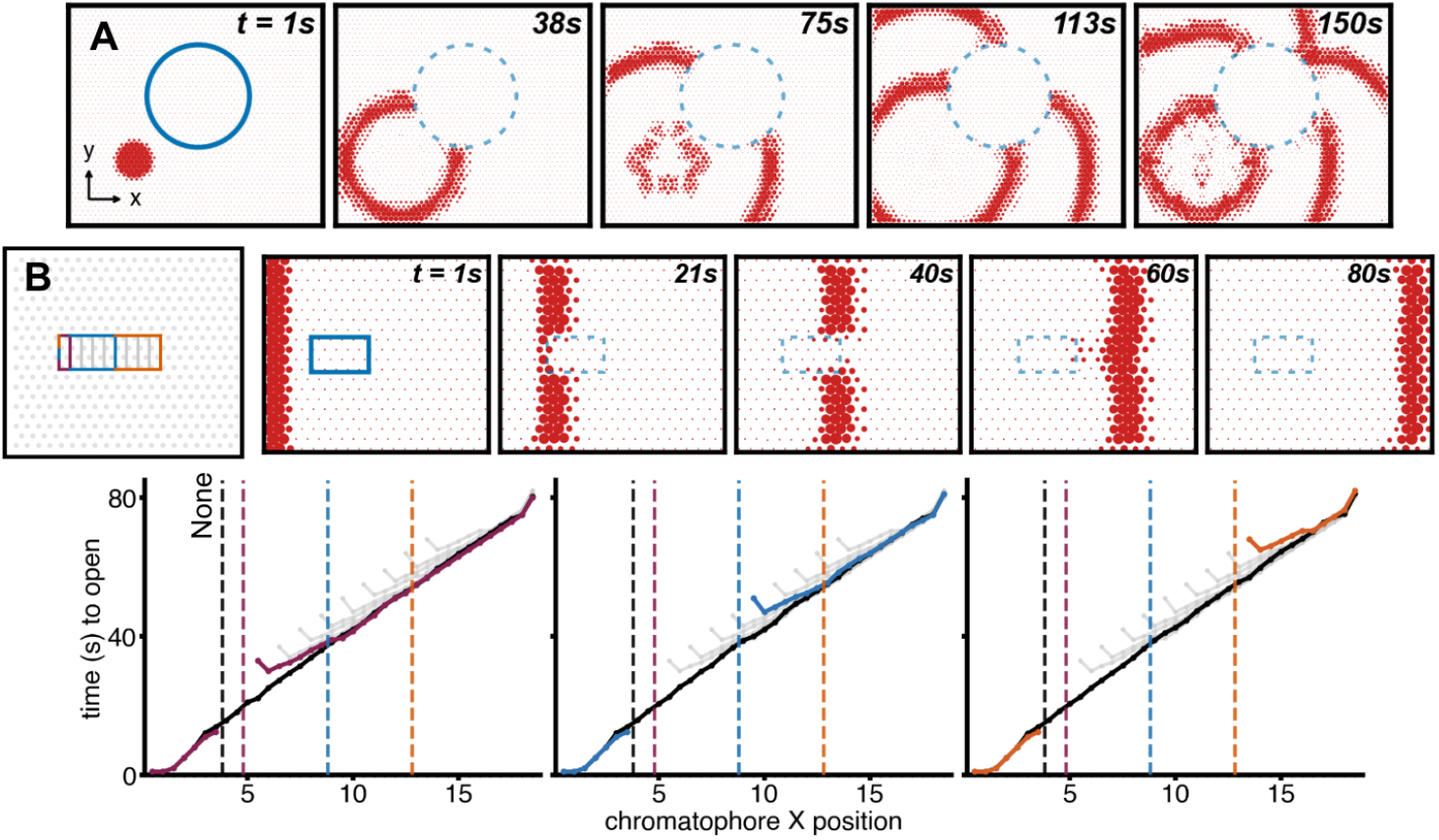
Rigid regions in the skin redirect, but do not block, traveling waves. (A) Circular obstacle. A transient point stimulus is applied near the lower-left corner of a 20 ×20 hexagonal network containing a rigid circular region in the upper right. Snapshots show the wave propagating around the obstacle; the two arms of the wave reunite behind the rigid region and continue as a reentrant wave. (B) Rectangular obstacles of increasing length. Upper left: schematic of the 20 ×20 network with rectangular rigid regions of length 1 (purple), 5 (blue), and 9 (orange) (grey indicates other intermediate lengths that were also tested). Upper right: representative snapshots from wave propagation around the blue rigid object. Lower: Wave arrival time (first opening time for chromatophores inside the *y*-position range of the rigid object) plotted against *x*-position for all three obstacle widths. Black line: baseline (no obstacle). Colored dashed vertical lines mark the end of each rigid band. Within the obstacle, chromatophores remain retracted as the wave routes around the obstruction; beyond the obstacle the slope returns to baseline, indicating that propagation speed has fully recovered. The offset between colored and black curves quantifies the total delay imposed by each obstacle, as a function of its size.

We examine various shapes and sizes for rigid objects to quantify how a passing wave is affected by chromatophores that do not open. The edge-initiated traveling plane wave is able to pass rectangular rigid bands of varying widths (Fig 6B; S10–S14 Videos). Within each rectangle (band), chromatophores remain unchanged as the wave routes around the band; beyond the band, the wave’s arrival-time profile resumes the slope of the no-obstacle baseline, showing that propagation speed fully recovers after the obstruction. The recovered section of the wave catches up quickly. The return delay scales relative to obstacle width, but the reconstitution is consistent in every case, even at the largest size, where the rigid band occupies nearly half the travel distance (Fig 6B, bottom right). This mimics the relative size of a large mantle patch that appears in some forms of disruptive coloration in cephalopods. ChromatoNet’s robustness in maintaining wave coherence around regions of static skin patterning mirrors observations in cuttlefish [4, 19].

### Varying connectivity

The sizes, numbers of colors, density per patch of skin, and connectivity among the chromatophores vary across cephalopods. The prevalence of inter-chromatophore muscle contacts are still being explored and the density of these contacts likely changes with developmental stage and may also vary with how the skin is dissected during excise experiments [35, 38]. The wave behaviors simulated above each assume a fully connected hexagonal lattice. We tested how robust those behaviors can be to partial connectivity.

Removing a fraction of randomly chosen muscle connections degrades wave propagation in stages. At low removal rates, waves initiated by an electrical pulse along the left edge still cross the network as a coherent front (Fig 7A, left). By 20% removal, waves typically reach the far edge but the wavefront is visibly fragmented, with isolated expanded chromatophores lagging or leading the bulk of the wave (Fig 7A, center). By 50% removal, the wave disperses into isolated, irregular clusters that rarely bridge the full network width (Fig 7A, right). At very high removal fractions (*>*75%), few chromatophores open at all; waves are rare and dispersion is ill-defined.

**Fig 7.**
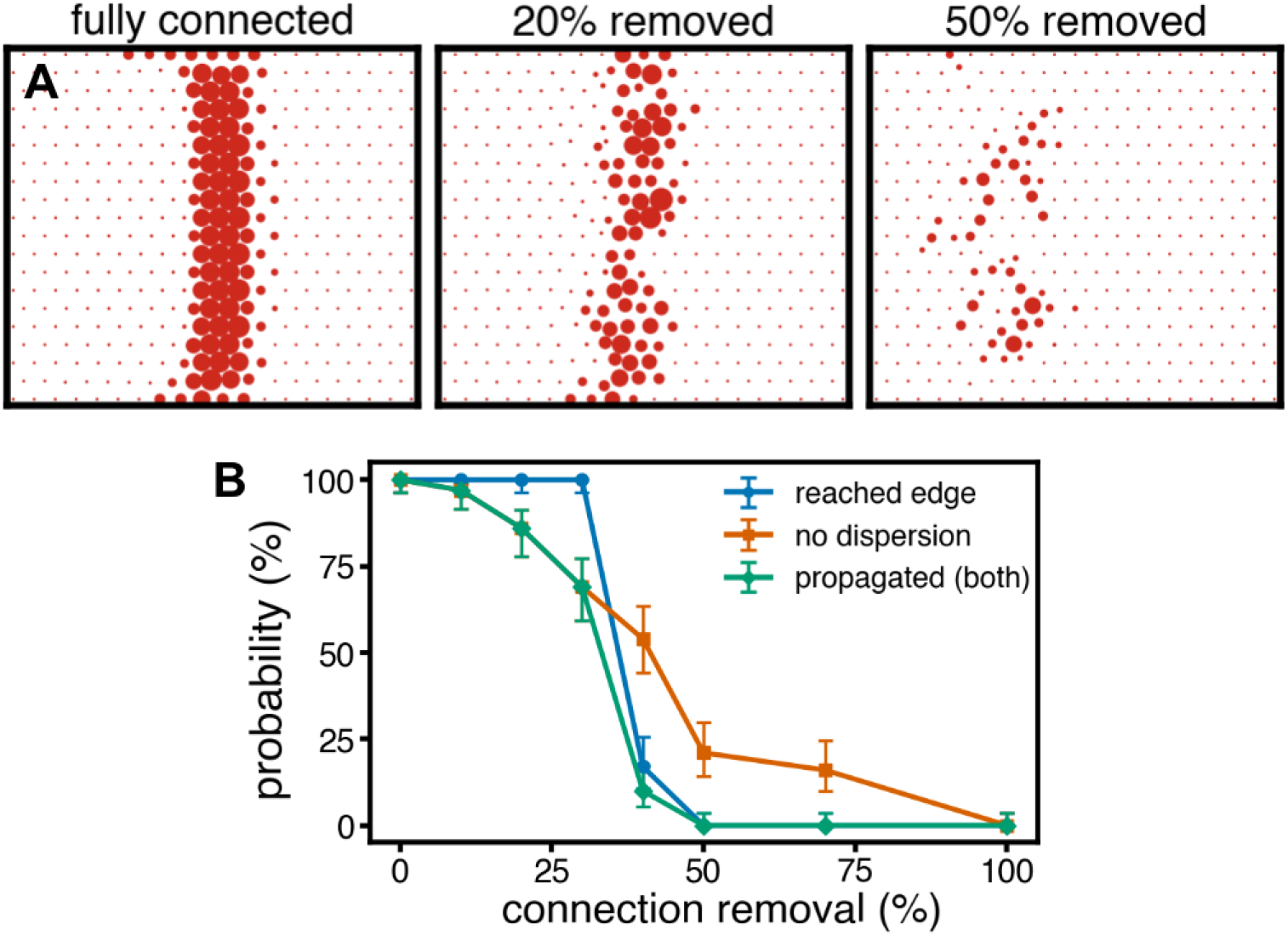
Network connectivity controls wave robustness and coherence. (A) Snapshots of a left-edge-initiated wave midway through propagation (approximately *t* = 40 s) for networks with decimation of 0% (fully connected, left), 20% (center), and 50% (right), wherein connections are removed at random, uniformly across the network. Sparse networks produce fragmented, irregular wave fronts; at 50% decimation, coherent propagation across the network is rare. (B) Quantifying propagation across 100 independent trials per decimation fraction. Blue: probability that majority of chromatophores on the right edge opened (wave reached the far edge); orange: probability that the wave remained spatially coherent (no dispersion into isolated or random sparse patches); green: probability that both criteria were satisfied (the wave propagated across as a unit). All curves are shown with ±1 SD error bars.

Quantifying this trend across 100 trials per removal fraction (Fig 7B) reveals a sharp transition. Both the probability of a wave reaching the far edge and the probability of remaining spatially coherent drop steeply between 30% and 50% removal, and the joint probability of satisfying both criteria falls to near zero by 50% removal. The behavior is consistent with percolation-like dynamics in sparse random networks [43]: there is a critical connectivity below which long-range propagation fails, and this threshold is robust across random-removal realizations (narrow error bars at intermediate decimation fractions).

The percolation threshold gives biology room to maneuver: coherent waves propagating in denervated cephalopod skin [35, 38] imply in-vivo networks sit above this threshold, though the ranges of precise connectivity fraction across species remain unmeasured.

## Discussion

We have shown how a simulation (ChromatoNet) of cephalopod skin composed of chromatophores and muscle connections with active response processes can imitate patterns that real cephalopods display. ChromatoNet’s elementary peripheral coupling rules, parameters, and muscle dynamics are sufficient to generate a diverse repertoire of patterns that is similar to those seen in living and denervated cephalopod skin. This includes spontaneous and evoked traveling waves, flickering, spirals, and body-wide flashes. Simple inputs can also produce waves that move around barriers. The wave speed and intrinsic frequency behaviors of the model match phenomena noted in real squid skin, without adjusting the model to specifically match each of those observations. This diverse output capacity suggests that the cephalopod brain may offload some aspects of its dynamic skin patterning mechanisms to locally autonomous, yet centrally controllable, effectors in the skin.

### ChromatoNet joined with experimental data

Several future routes could align ChromatoNet more firmly to live cephalopod data. Existing imaging tools are already available to quantify skin features more fully, and the experiments and analyses suggested below would close the loop between this model and the animals it represents.

#### Wet-lab tests of model predictions

Several model predictions are testable in a denervated skin preparation. Bath application of pharmacological agents targeting gap junctions, voltage-gated calcium channels, or membrane conductances would shift 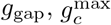 or *g*_*L*_, each with predicted consequences for wave speed and frequency.

Stimulating denervated skin under each manipulation and measuring the resulting traveling-wave speed or focal-wave frequency would provide a direct quantitative test. Pharmacological agents could modulate effective skin elasticity and hence inform model spring constants. A separate prediction is that spontaneous flickering should peak at an intrinsic frequency, accessible by power-spectrum measurements of videos of live animals. Both experiments are within reach of established techniques [35, 38].

Direct mechanical measurements of cephalopod muscles would also tighten the model. The mechanical parameters (*k*_*s*_, *k*_*c*_, *k*_*p*_, *L*_0_) have not been measured experimentally, and the values used here produce biologically plausible wave speeds but should be treated as order-of-magnitude estimates until such measurements become available. Quantifying the actual sources of stochasticity in real chromatophore muscles, which may include spontaneous calcium release events [37], would similarly sharpen the noise model used for spontaneous-activity simulations.

#### Brain-stimulation experiments and time-evolving displays

Boycott et al. [2] and Reiter et al. [32] mapped chromatophore-lobe stimulation to skin regions in chronic cuttlefish recordings, providing a direct link between central commands and peripheral activation, as did earlier work in squid (Dubas et al. [7]) chromatophore lobe. Using these stimulus-response maps as inputs to ChromatoNet would let the model predict what happens to an activated region after the central stimulus releases: does it fade passively, propagate outward as a wave, or seed a sustained dynamic pattern? Comparing predictions to the in vivo time evolution would test whether the peripheral substrate alone accounts for post-stimulation dynamics, or whether ongoing central drive is needed.

#### Anatomically grounded networks from video

ChromatoNet could be used to fit videos of live cephalopod skin and suggest what biophysical parameters create the observed patterns *in vivo*. Existing tools extract chromatophore positions from video [32, 42], making it feasible to construct a ChromatoNet network directly from a sequence of frames in two complementary modes. First, chromatophore positions can be extracted from the video and used directly as lattice positions for the model, with muscles assigned to the *N* nearest neighbors. Second, both positions and color class can be extracted, with muscle weights assigned based on same-color proximity (closer same-color chromatophores get higher weights) and on layer depth (cross-layer attachments scaled by distance through the depth axis). Current pipelines such as CHROMAS [42] handle position tracking but color segmentation often requires manual annotation. Improved color-class detection is itself a useful direction for the field. Video data also carry dynamics, not just anatomy. A sufficiently long recording could constrain the model’s intrinsic parameters: extract chromatophore time courses, fit ChromatoNet parameters to reproduce those dynamics, and read off biophysical constants directly from observed cephalopod behavior.

Denervated and electrically stimulated preparations are the ideal candidates for this fitting, since they remove the brain signal as a hidden variable. A further opportunity along these lines is longitudinal imaging of live animals, in which a young cephalopod is imaged repeatedly throughout development. Tracking the same chromatophores over time would reveal which new chromatophores appear, which existing ones grow, and (if spatial resolution allows) how their muscles extend and connect to neighbors, providing direct experimental input for developmental network growth simulations.

### Extending the model: geometry and color

Currently, the default ChromatoNet simulation works on a single layer of monochromatic chromatophores connected in a regular hexagonal lattice, with uniform parameters across the network. Real cephalopod skin departs from this in several ways that current results do not yet address. For example, squids typically have a superficial layer of yellow chromatophores, and a middle layer of reds and a basement layer of browns [11]. Specific extensions can move the model closer to the underlying anatomy.

#### Connectivity radius and muscle range

A first step toward more realistic model geometry is to extend the connectivity radius beyond nearest neighbors, since each chromatophore is surrounded by 20–30 radial muscles in real squid skin that can extend to more distal neighbors [5, 20, 22, 38]. We ran simulations in which each chromatophore connects to its first (6), second (6), and fourth (12) neighbor shells in a hexagonal setup (24 muscles per chromatophore; S2 Fig). The chromatophores at the third-neighbor distance were skipped since they are radially aligned to the first-neighbor connections. Wave behavior in this 24-connect network was qualitatively similar to the 6-connect case across the parameter regimes tested (S15 Video). The basic dynamics are robust to changes in the connectivity radius.

Senft et al. [38] report that some chromatophore muscles in *Doryteuthis pealeii* project beyond their immediate neighbors, consistent with the picture that real skin contains a mix of short- and longer-range connections. A natural extension would sample muscle reach from a random or generated distribution rather than fixed layers, allowing the network to capture the heterogeneous connectivity likely present *in vivo*.

#### Variability across chromatophores and muscles

The current model assigns identical parameters to every chromatophore and every muscle, but real skin is heterogeneous. For example, in squid skin, yellow chromatophores are smallest, reds next and browns largest [5, 11, 22]. Chromatophore size and number also varies systematically with developmental age, with older chromatophores larger than younger ones that intercalate into the arrays over time [28]. Recent quantitative imaging of squid mantles confirms that the size distribution is broad and stationary during growth [36], with larger older chromatophores surrounded by smaller/younger ones in a disordered arrangement.

Muscle strength similarly may vary across cells, though this has not been quantitatively measured. Anatomical work has identified different morphological types of chromatophore muscles [38], but strength differences across types have not been characterized.

Another extension would draw chromatophore size, muscle stiffness, and other single-element parameters from distributions, which can reveal whether the wave behaviors identified here are robust to plausible biological heterogeneity or specifically require well-matched elements.

#### Tissue anchoring as discrete connections

The current model assumes every chromatophore is anchored to tissue with the same skin spring constant *k*_*s*_. Real skin includes muscles that connect chromatophores not only to each other but also directly to the surrounding tissue (Fig 1B) [38]. Treating muscle-to-tissue anchoring as springs that vary across chromatophores would represent localized stiffening more naturally. This would allow us to use the model to explore how gap junction dynamics differ between muscle-to-muscle and muscle-to-tissue contacts (since gap junctions are modeled as forming between any neighboring muscles regardless of their attachment partner), and to test the effects of connection-removal in muscle-to-tissue contacts as well as muscle-to-muscle ones.

Senft et al. [38] document that these connections exist but do not fully characterize their spatial distribution, leaving room for more anatomical work to constrain the model. Additionally, it should be possible using antibodies to gap junctions and stretch channels to verify that these components are present where muscles overlap.

#### Layers, colors, and size

Real squid skin contains multiple layers of chromatophores in different color classes, and the spatial arrangement of those layers directly shapes which chromatophores are likely to connect to which others. A characteristic spatial signature is that a large, dark chromatophore is typically surrounded by smaller, lighter ones that occupy the spaces it left open [11, 22, 28], producing a scale-free arrangement of pigment elements (Fig 1A). Chromatophores arise continuously throughout the animal’s life and maintain the same morphological geometry. Pigment sacs start lighter and progressively darken as they migrate through the skin and age, so the oldest chromatophores are the largest and the darkest (brown). The colors live in successive layers of the skin, with the darkest deepest and the lightest most superficial. Chromatophores of the same color class are therefore also spatially proximate in depth.

To extend the model toward this anatomy, the network could be grown from a developmental rule rather than wired by hand. A simple version: (1) seed the deepest layer with a sparse set of large, dark chromatophores, then grow muscles from each chromatophore radially outward at randomized speeds, with muscles attaching to other muscles when they encounter them within a threshold distance; (2) add a second layer of smaller, lighter chromatophores between existing ones, growing their muscles into 3D space, with same-layer attachments favored by proximity but cross-layer attachments possible and weighted by distance through depth; (3) repeat with progressively newer, lighter generations, with pigment color darkening over the simulated lifespan. This would naturally produce denser within-color coupling and sparser cross-color coupling without imposing it by hand, and would let the model track the geometry of the skin of an animal from hatchling to adult.

## Conclusion

ChromatoNet simulations reveal a handful of global and local parameters, model “knobs” that might have biological analogs, allowing interpolation between qualitatively different behavioral regimes. This suggests that signals, perhaps neuromodulatory, might shift the skin between communication and camouflage modes without requiring direct control of each chromatophore. Cephalopod skin may not be a passive display medium waiting for instructions, but an excitable substrate with its own dynamics, tuned to produce rich patterns from minimal input. The brain may possess the flexibility to conduct the chromatophore orchestra rather than calculating every note.

## Supporting information

**Fig S1.**
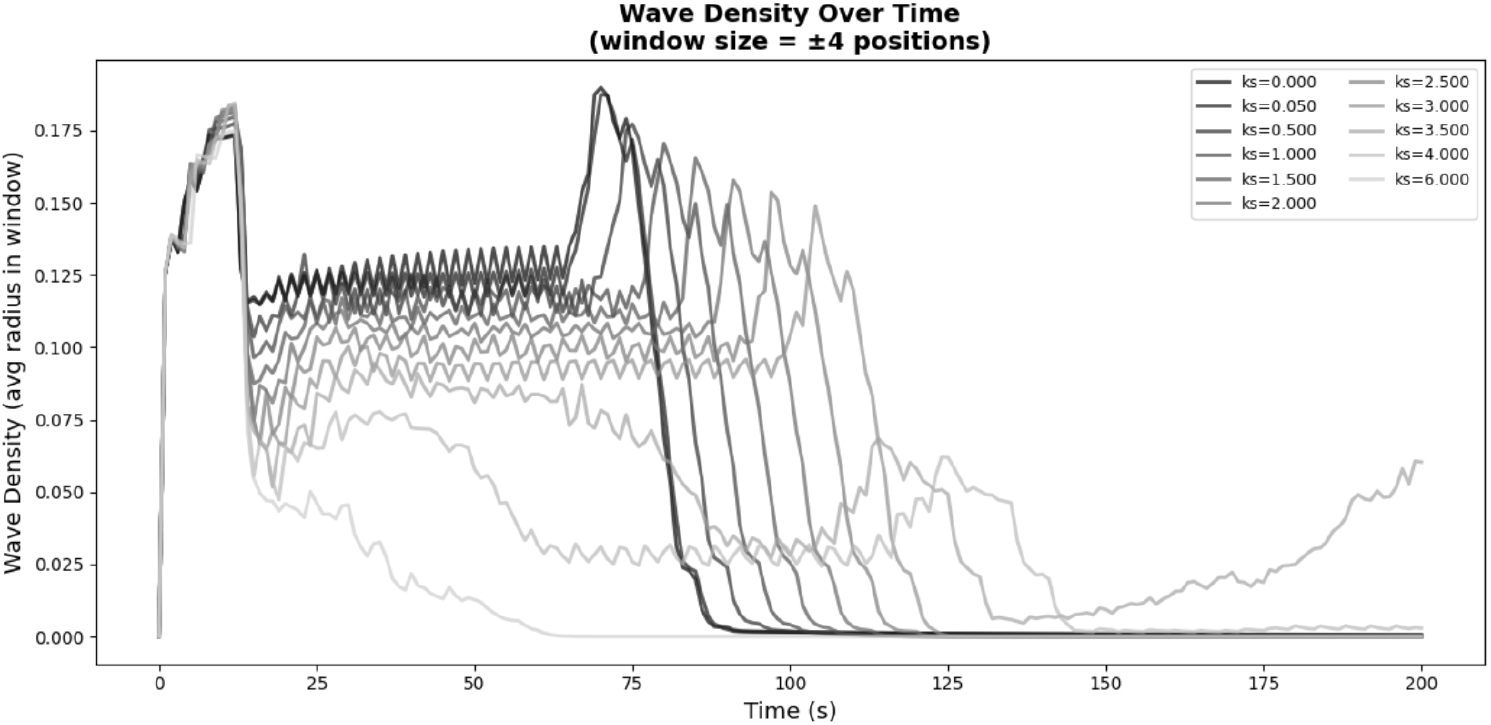
Wave density as a function of skin spring constant *k*_*s*_. Sample of parameter effect analysis: wave density across the *x*-axis for a sweep of *k*_*s*_ values, complementing Fig 5D.

**Fig S2.**
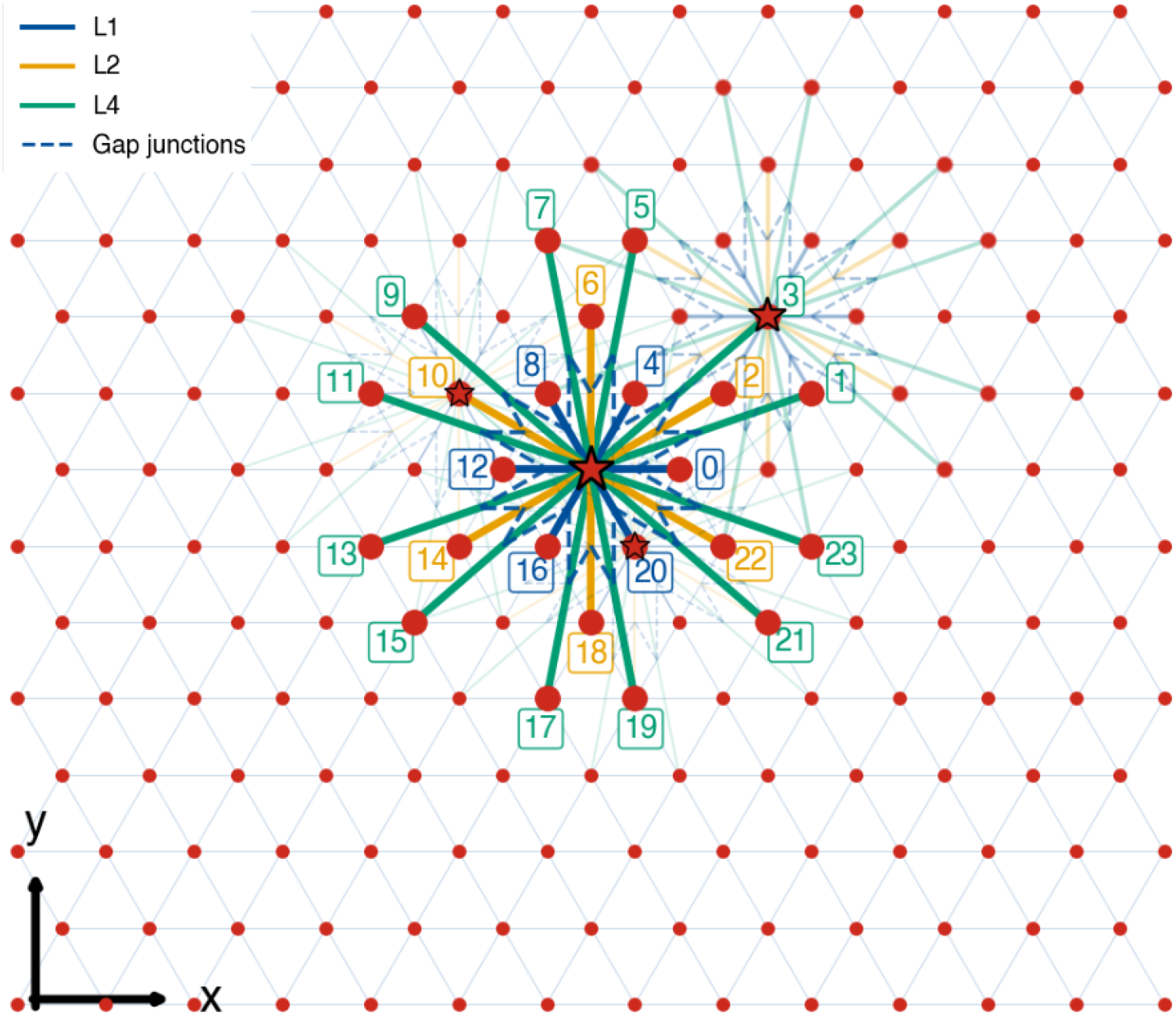
Extended connectivity (24-connect). Graph representation of the extended connectivity scheme: each chromatophore is connected to its first, second, and fourth neighbor shells (L1, L2, L4), for a total of 24 muscles per chromatophore, with gap junctions between neighboring muscles within each shell. At the parameter regimes tested in this paper, wave behavior with 24-connect coupling is qualitatively similar to the 6-connect case used in the main figures.

**S1 Video. Traveling waves in a flamboyant cuttlefish**. A flamboyant cuttlefish (*Ascarosepion pfefferi*) feeding while dark traveling waves pass across its mantle, including through regions of skin that hold a static pattern. Video courtesy of the Hanlon Laboratory.

**S2 Video. Traveling wave across a squid mantle**. A traveling wave crossing the mantle of *Doryteuthis pealeii* ; source footage for Fig 2B. Video courtesy of the Hanlon Laboratory.

**S3 Video. Traveling wave in the 1D model**. Animation corresponding to Fig 2A. A brief positional displacement of the leftmost chromatophore initiates a wave that propagates along the chain.

**S4 Video. Traveling wave in a** 50 × 50 **hexagonal network**. Traveling wave on a 50 × 50 hexagonal lattice with *g*_*L*_ = 0.4; all other parameters as in Table 1.

**S5 Video. Spiral wave**. Animation corresponding to Fig 2C (bottom). A transient asymmetric stimulus near a rigid strip splits the wavefront; the two arms propagate around the obstacle and reenter to form a persistent rotating spiral.

**S6 Video. Periodic reentrant wave from an asymmetric stimulus**. Animation corresponding to Fig 3A (bottom). A transient stimulus restricted to a narrow strip of the edge produces a wave that reflects and reenters, generating a periodic traveling wave from a single input.

**S7 Video. Spontaneous activity in excised cephalopod skin**. Continuous spontaneous chromatophore activity in pinned, excised skin of *Ascarosepion pfefferi* ; source footage for Fig 4C. Video courtesy of the Hanlon Laboratory.

**S8 Video. Spontaneous noise-driven activity in the model**. Animation corresponding to Fig 4B (top). Frozen Gaussian noise (*σ* = [TO FILL]) added to chromatophore rest positions produces wave fragments that nucleate, propagate, and dissolve without external stimulation.

**S9 Video. Reentrant wave around a circular rigid region**. Animation corresponding to Fig 6A. A transient stimulus near the lower left of a 20 × 20 network splits around a rigid circular region in the upper right; the arms reunite behind it and establish a periodic reentrant wave.

**S10 Video. Traveling wave passing a rigid band of length 1**.

Left-edge-initiated plane wave on a 20× 20 network routing around a rectangular rigid region of length 1 (Fig 6B).

**S11 Video. Traveling wave passing a rigid band of length 3**. As in S10 Video, with a rigid region of length 3.

**S12 Video. Traveling wave passing a rigid band of length 5**. As in S10 Video, with a rigid region of length 5 (blue in Fig 6B).

**S13 Video. Traveling wave passing a rigid band of length 7**. As in S10 Video, with a rigid region of length 7.

**S14 Video. Traveling wave passing a rigid band of length 9**. As in S10 Video, with a rigid region of length 9 (orange in Fig 6B).

**S15 Video. Traveling wave in a 24-connect network**. Animation of a left-edge-initiated traveling wave on a hexagonal lattice with extended connectivity (L1, L2, L4 shells; 24 muscles per chromatophore), corresponding to the connectivity schematic in S2 Fig.

**S16 Video. Direct stimulation of a rigid region with unpinned edge boundaries**. Animation of a 20×20 hexagonal network containing a rigid stripe region in which the edge chromatophores are not held fixed at their rest positions, unlike in all other simulations. A transient stimulus applied directly within the rigid region produces the activity shown: activated rigid region with reentrant traveling waves.

## Acknowledgments

This work was supported by the Polymaths Program from Schmidt Sciences, LLC (SEP). This work was supported by the National Science Foundation through the Physics Frontier Center for Living Systems (PHY-2317138). This work was supported by the NSF-Simons National Institute for Theory and Mathematics in Biology, awards NSF DMS-2235451 and Simons Foundation MP-TMPS-00005320. We acknowledge partial support from Grant FA9550-22-1-0467 from the Air Force Office of Scientific Research. We thank Zach Morrissey for his support and for sharing colorblind-friendly figure design resources.

## Notes

### Competing Interest Statement

The authors have declared no competing interest.

