## Supplementary figures and images for "Bio-inspired chromatophore network model generates dynamic skin patterns akin to those observed in cephalopods"

### Supplemental Figure 1

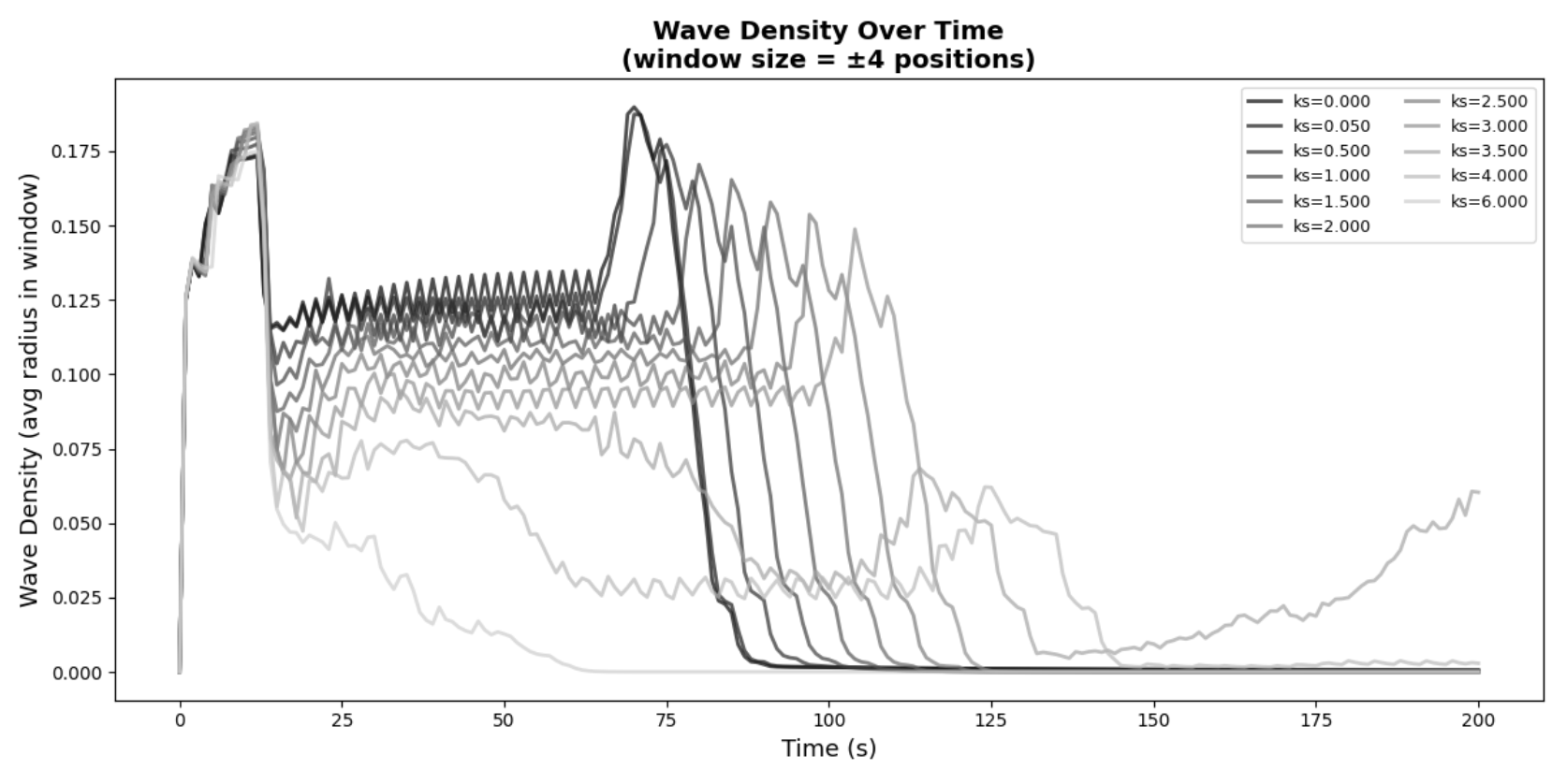

### Supplemental Figure 2

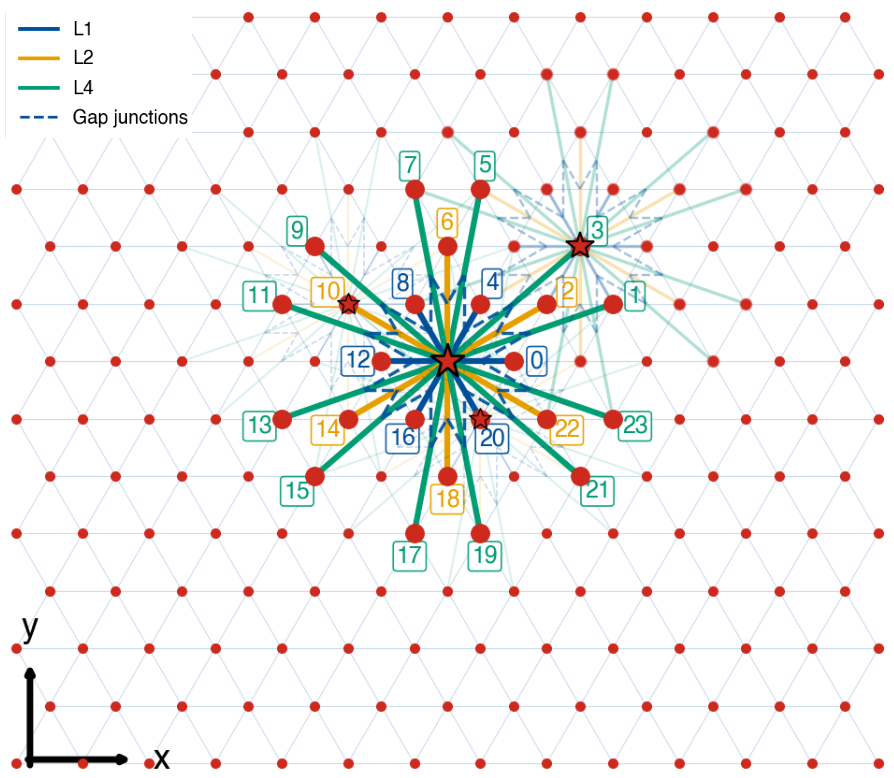

### Supplemental Video 2

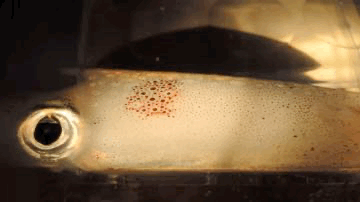

### Supplemental Video 3

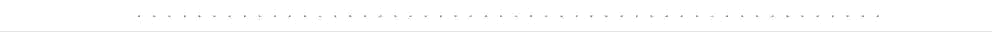

### Supplemental Video 15

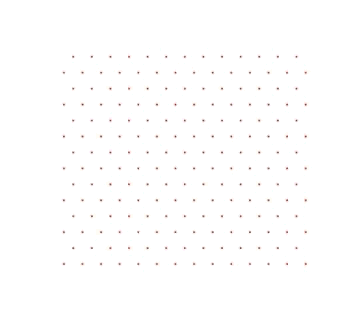
